# Chitinase 3-like-1 (CHI3L1) Inhibits Innate Anti-Tumor and Tissue Remodeling Immune Responses by Regulating CD47-SIRPα and CD24-Siglec10-Mediated Phagocytosis

**DOI:** 10.1101/2023.12.27.573478

**Authors:** Bing Ma, Suchitra Kamle, Chang-Min Lee, Joyce H Lee, Daniel C Yee, Zhou Zhu, Edwin K. Silverman, Dawn L. DeMeo, Augustine M.K. Choi, Chun Geun Lee, Jack A. Elias

## Abstract

Innate immune responses such as phagocytosis are critically linked to the generation of adaptive immune responses against the neoantigens in cancer and the efferocytosis that is essential for homeostasis in diseases characterized by lung injury, inflammation, and remodeling as in Chronic Obstructive Pulmonary Disease (COPD). Chitinase 3-like-1 (CHI3L1) is induced in many cancers where it inhibits adaptive immune responses by stimulating immune checkpoint molecules (ICPs) and portends a poor prognosis. CHI3L1 is also induced in COPD where it regulates epithelial cell death. Here we demonstrate that pulmonary melanoma metastasis inhibits macrophage phagocytosis by stimulating the CD47-SIRPα and CD24-Siglec10 phagocytosis checkpoint pathways while inhibiting macrophage “eat me” signals from calreticulin and HMGB1. We also demonstrate that these effects on macrophage phagocytosis are mediated by CHI3L1 stimulation of the SHP-1 and SHP-2 phosphatases and the inhibition of the accumulation and phosphorylation of cytoskeleton-regulating non-muscle myosin IIa. This inhibition of innate immune responses like phagocytosis provides a mechanistic explanation for the ability of CHI3L1 to stimulate ICPs and inhibit adaptive immune responses in cancer and diseases like COPD. The ability of CHI3L1 to simultaneously inhibit innate immune responses, stimulate ICPs, inhibit T cell co-stimulation, and regulate a number of other oncogenic and inflammation pathways suggest that CHI3L1-targeted therapeutics are promising interventions in cancer, COPD and other disorders.

## Introduction

Innate and adaptive immunity are dysregulated in and play important roles in the pathogenesis of inflammatory, remodeling, neoplastic and other diseases and disorders (1). In cancer, neoantigens are generated and recognized as non-self, allowing the host to mount T-cell mediated immune responses that can control and/or kill the tumor cells (2). Tumor initiation and or progression are frequently seen when these protective adaptive immune responses are inhibited by the induction of immune checkpoint molecules (ICPs) as in the PD-1-PD-L1/L2, CTLA4-B7.1/B7.2 and LAG3 pathways (3, 4). Recent studies have demonstrated that the onset and maintenance of these immunosuppressive immune responses and the development of long-lasting memory T cells requires innate immune activation (5, 6). Specifically, cells such as innate immune macrophages launch antigen-specific adaptive responses while mounting their own effector responses such as phagocytosis (5, 6). This is particularly important in neoplasia where mononuclear phagocyte ingestion of tumor cells leads to the cross presentation of tumor neoantigens by MHC class I on the cell surface which, in turn, induces the priming and accumulation of tumor suppressing antigen-specific CD8+ T cells (5). Although phagocytosis is believed to be a critical modality that serves as a bridge between innate and adaptive immunity (7), the mechanisms that activate innate immune responses in cancer and other disorders have only recently begun to be appreciated.

Phagocytosis of apoptotic cells, termed efferocytosis, is paramount for the maintenance of tissue homeostasis (7). This is due to the ability of phagocytosis to prevent secondary necrosis of apoptotic cells and the release of autoantigens and proinflammatory alarmins (7). These anti-inflammatory effects are particularly important at mucosal locations such as the lung where the epithelium can turn over every 7-21 days (7–13). Efferocytosis is an evolutionarily conserved, highly regulated, multistep process that includes mononuclear cell recognition of apoptotic cells and the proper interpretation of the signals they send (7). Specifically, apoptotic cells display diverse “don’t eat me” signals such as CD47, CD24, CD31 and PD-L1 and “eat me” signals such as calreticulin, HMGB1 and annexin-1 (7, 14, 15). The ultimate effect of these signals is additive and depends on the balance of phagocytosis inhibiting and stimulating signals and their respective ligands (7). This balance plays an important role in malignancies which try to evade phagocytosis by stimulating anti-phagocytotic “don’t eat me” and inhibiting “eat me” receptors and their ligands (7). Efferocytosis also plays a key role in inflammatory and remodeling disorders like COPD which is characterized by Th1/Th17 inflammation, neutrophil accumulation, the accumulation of uncleared apoptotic cells that can undergo secondary necrosis and deficiencies in efferocytosis. (16–22). However, the pathways that regulate phagocytosis in these and other settings have not been adequately defined.

Chitinase 3-like-1 (*CHI3L1*; also called *Chil1* (mouse) and *YKL-40* (human)) is a member of the 18 glycosyl hydrolase gene family that is produced by myeloid, epithelial, endothelial and other immune cells (23–30). Studies from our laboratory and others have highlighted its pleiotropic effects including its ability to inhibit apoptosis, stimulate M2 macrophage activation and differentiation, inhibit oxidant injury and stimulate fibroproliferative healing and repair (23, 24, 26, 31). They also described the receptors that mediate many of these and other responses including the multimeric chitosome which has IL-13Rα2 as its alpha subunit and TMEM219 or CD44 as its beta subunits (27, 32, 33) and CRTH2 (26). CHI3L1-receptor binding can induce MAP Kinase, Akt/PKB, and or Wnt/μ-catenin signaling (27, 34). CHI3L1 is readily detected in the circulation of normals and elevated levels of circulating CHI3L1 are seen in patients with inflammatory, remodeling and aging related disorders (25, 35–38). The levels of circulating CHI3L1 are also increased in patients with visceral cancers where they correlate with disease progression and inversely with disease free interval and survival (3, 39–46). Recent studies from our laboratory have also demonstrated that CHI3L1 is also a master inducer of antitumor adaptive immune responses including the activation of the PD-1, PD-L1, PD-L2; CTLA4, B7.1, B7.2; and LAG 3 pathways and an inhibitor of T cell costimulation via Icos, IcosL and CD-28 (3, 4). Studies from our laboratory and others have demonstrated that CHI3L1 is also induced by cigarette smoke exposure where it inhibits epithelial injury and emphysema generation and is associated with dysfunctional adaptive immune responses (37, 47). Although CHI3L1 is induced in the circulation of patients with cancer and remodeling disorders like COPD and is a potent regulator of cellular apoptosis, the ability of CHI3L1 to regulate innate immune responses such as phagocytosis and the mechanisms that CHI3L1 uses in these regulatory events have not been defined.

We hypothesized that CHI3L1 is a critical regulator and activator of innate immune responses like phagocytosis and that this activation plays a critical role in the ability of CHI3L1 to induce ICPs and inhibit adaptive immune responses. To test this hypothesis, we used *in vitro* and *in vivo* approaches to characterize the ability of CHI3L1 to regulate phagocytosis by regulating “don’t eat me” and “eat me” signals and characterizing the roles of non-muscle myosin IIa (NM IIa) in mediating these responses. These studies demonstrate that CHI3L1 is a powerful inhibitor of phagocytosis that mediates its effects by stimulating the CD47-SIRPα and CD24-Siglec10 “don’t eat me” and inhibiting the calreticulin and high mobility group box protein −1 (HMGB-1) “eat me” phagocytosis pathway signals. These events inhibited the phosphorylation and disrupted NM IIa which is the major regulator of the cytoskeletal rearrangements that occur during phagocytosis (14). These studies demonstrate that CHI3L1 regulates adaptive antitumor responses and stimulates ICPs via the regulation of innate immunity and phagocytosis via modulating “don’t eat me” and “eat me” innate immune signals and their ligands. They also support the potential utility of inhibitors of CHI3L1 as multifaceted novel inhibitors of oncogenesis and tissue inflammation and remodeling.

## Materials and Methods

### Genetically modified mice

Mice with null mutations of Chil1 *(Chil1^-/-^)* and transgenic mice in which *CHI3L1* was targeted to the lung with the CC10 promoter (*CHI3L1 Tg*) have been generated and characterized by our laboratory as previously described (23, 27). These mice were between 6-8 weeks old when used in these studies. All animals were humanely anesthetized with Ketamine/Xylazine (100mg/10mg/kg) before any intervention. The protocols that were used in these studies were evaluated and approved by the Institutional Animal Care and Use Committee (IACUC) at Brown University.

### Western blot analysis

Protein lysates from macrophages and whole mouse lungs were prepared with RIPA lysis buffer (ThermoFisher Scientific, Waltham, MA, USA) containing protease inhibitor cocktail (ThermoFisher Scientific) as per the manufacturer’s instructions. 20 to 30 µg of lysate protein was subjected to electrophoresis on a 4–20% gradient mini-Protean TGX gel (Bio-rad, Hercules, CA, USA). It was then transferred to a PVDF membrane using a semidry method with a Trans-Blot Turbo Transfer System (Bio-rad). Membranes were blocked with Tris-buffered saline with Tween20 (TBST) with 5% non-fat milk for 1 hour at room temperature. After blocking, the membranes were incubated with the primary antibodies against CD47 (Invitrogen, miap301), SIRPα (Invitrogen, PAS-97176), CD24 (eBioscience, M1/69), Siglec10 (R&D Systems, MAB7775), HMGB1 (Abcam, ab-79823), calreticulin (Invitrogen, 1G6A7); SHP-1 (Cell Signaling Technology, C14H6), and SHP-2 (Cell Signaling Technology, #3752) overnight at 4°C in TBST and 5% BSA. The membranes were then washed 3 times with TBST and incubated with secondary antibodies in TBST, 5% non-fat milk for 1 hour at room temperature. After 3 additional TBST washes, Supersignal West Femto Maximum Sensitivity Substrate Solution (ThermoFisher Scientific) was added to the membrane and immunoreactive bands were detected by using a ChemiDoc (Bio-Rad) imaging system. ImageJ was used to quantitate the relative signal intensity of each Western blot.

### RNA extraction and Real-time qPCR

Total cellular RNA was obtained using TRIzol reagent (ThermoFisher Scientific) followed by RNA extraction using RNeasy Mini Kit (Qiagen, Germantown, MD) according to the manufacturer’s instructions. mRNA was measured and used for real time (qRT)-PCR as described previously (23, 31). The primer sequences used in these studies are summarized in Table S1. Ct values of the test genes were normalized to the internal housekeeping gene β-actin.

### Immunohistochemistry

Formalin-fixed paraffin embedded (FFPE) lung tissue blocks were serially sectioned at 5 μm-thickness and mounted on glass slides. After deparaffinization and dehydration, heat-induced epitope retrieval was performed by boiling the samples in a steamer for 30 minutes in antigen unmasking solution (Abcam, antigen retrieval buffer, 100X citrate buffer pH 6.0). To prevent nonspecific protein binding, all sections were blocked in a ready-to-use serum free protein blocking solution (Dako/Agilent, Santa Clara, CA) for 10 minutes at room temperature. The sections were incubated with primary antibodies against CD47(Invitrogen, miap301), SIRPα, (Invitrogen, PAS-97176), CD24 (eBioscience, M1/69), and Siglec10 (R&D Systems, MAB7775) overnight at 4°C. After three washings, the slides were incubated for 1 hour at room temperature with fluorescence labeled secondary antibodies. The sections were then mounted with mounting medium, stained with DAPI dye and images were acquired using a fluorescent microscope (Nikon Ti2).

### Assessment of macrophage Phagocytosis

The mouse macrophage RAW 264.7 cell line was purchased from ATCC (Cat# TIB-71). The cells were cultured to confluence in RPMI medium with 10% FBS and 1% penicillin and streptomycin, inoculated into four chamber slides at a concentration of 2.0X10^5^/ml at 37°C and incubated overnight. The next day the cells were placed in RPMI medium without FBS for 4 hours. They were then incubated with recombinant (r) CHI3L1 (500ng/ml, R&D Systems, #2599-CH) in the presence of anti-CHI3L1 antibody (FRG, 250ng/ml) or with IgG2b isotype control antibody or the chitinase inhibitor kasugamycin (250ng/ml) for 48h. Zymosan bioparticles conjugated with flurescence-488 (2.0X10^5^/ml, Invitrogen, Z23373) were then added to each well and incubated for 30 min. At the end of this incubation, the slides were washed with PBS three times and the cells were stained with phalloidin (Invitrogen, 1:3000) for 10 min. After fixation of the slides with cold 4% paraformaldehyde for 10 min, fluorescence images were obtained using a. fluorescent microscope (Nikon, Ti2). In a similar way, pHrodo deep red E. coli bioparticles (2.0X10^5^/ml, Invitrogen, P35361), another advanced phagocytosis indicator, was also used to detect phagocytic macrophages. In addition to RAW cells, macrophages derived from THP-1 human monocytic cells were used to test the effect of CHI3L1 in phagocytic activities. In brief, THP-1 cells were stimulated with Phorbol Myristic Acetate (PMA, 25ng/ml) for 24 hours. They were then incubated with recombinant M-CSF (50ng/ml) and rCHI3L1 (R&D Systems, 250ng/ml) alone or together, in the presence of FRG (250ng/ml), kasugamycin (250ng/ml, Sigma, #32354) or their respective controls for 48 hours. The phagocytic activity of these cells was assessed using zymosan bioparticles as described above.

### Evaluation of Myosin IIa (NM2a)

The effect of CHI3L1 on NMIIa during macrophage phagocytosis was evaluated using RAW cells incubated with rCHI3L1 (500ng/ml). These cells were randomized to be incubated with antiCHI3L1 antibody (FRG, 250 ng/ml) or its IgG2b isotype (250ng/ml). After 30 minutes of incubation with zymosan bioparticles (1X10^6^/ml, Invitrogen), cells were harvested and subjected to the evaluation of NMIIa expression and activation (phosphorylation). Antibodies against total-myosin IIa (Cell Signaling Technology, #3752) and phosphorylated myosin IIa (Cell Signaling Technology, #3403) were used for immunoblotting analysis to detect the status of NMIIa.

### Fluorescence-activated cell sorting (FACS) analysis

The phagocytic activity of macrophages was also assessed using FACS analysis. RAW macrophage cells were cultured and challenged with rCHI3L1(500ng/ml) alone or together with FRG antibody (250ng/ml) or kasugamycin (250ng/ml) for 48 hours as described above. After 30-minute incubation with pHrodo deep red E. coli bioparticles (2X10^5^/ml, Invitrogen), the cells were fixed with 4% paraformaldehyde and subjected to FACS evaluation using the BD FACSAriaIIIu and FlowJo V10 software.

### Statistical Analysis

Statistical evaluations were undertaken with GraphPad Prism software. As appropriate, groups were compared with 2-tailed Student’s *t* test or with nonparametric Mann-Whitney *U* test. Values are expressed as mean ± SEM. One-way ANOVA or nonparametric Kruskal-Wallis test were used for multiple group comparisons. Statistical significance was defined as a level of *P* < 0.05.

## Results

### Pulmonary melanoma metastasis activates the CD47-SIRPα and CD24-Siglec10 “don’t eat me” phagocytic pathways via a CHI3L1-dependent mechanism(s)

Studies were undertaken to determine if major “don’t eat me” phagocytic pathways were regulated by pulmonary metastasis. In these experiments, wild type C57BL/6 mice (*Chil1*^+/+^) and *Chil1* null mutant mice (*Chil1*^-/-^) were challenged with freshly prepared B16-F10 melanoma cells (B16) or vehicle control and the levels of mRNA encoding CD47, SIRPα, CD24 and Siglec10 were assessed 2 weeks later. These studies demonstrated that B16 metastasis increased the levels of expression of CD47 and its ligand SIRPα and CD24 and its ligand Siglec10 (Figure 1, A-D). In all cases these inductive responses were dependent on CHI3L1 and were significantly decreased in comparisons of wild type (+/+) and *Chil1* null (-/-) mice (Figure 1, A-D). These alterations in mRNA accumulation were associated with comparable alterations in CD47, SIRPα, CD24, and Siglec10 protein accumulation as assessed by Western analysis (Figure 1E). These studies demonstrate that melanoma lung metastasis is associated with enhanced expression and induction of “don’t eat me” phagocytic signals and that this enhanced induction is mediated by CHI3L1.

**Figure 1.**
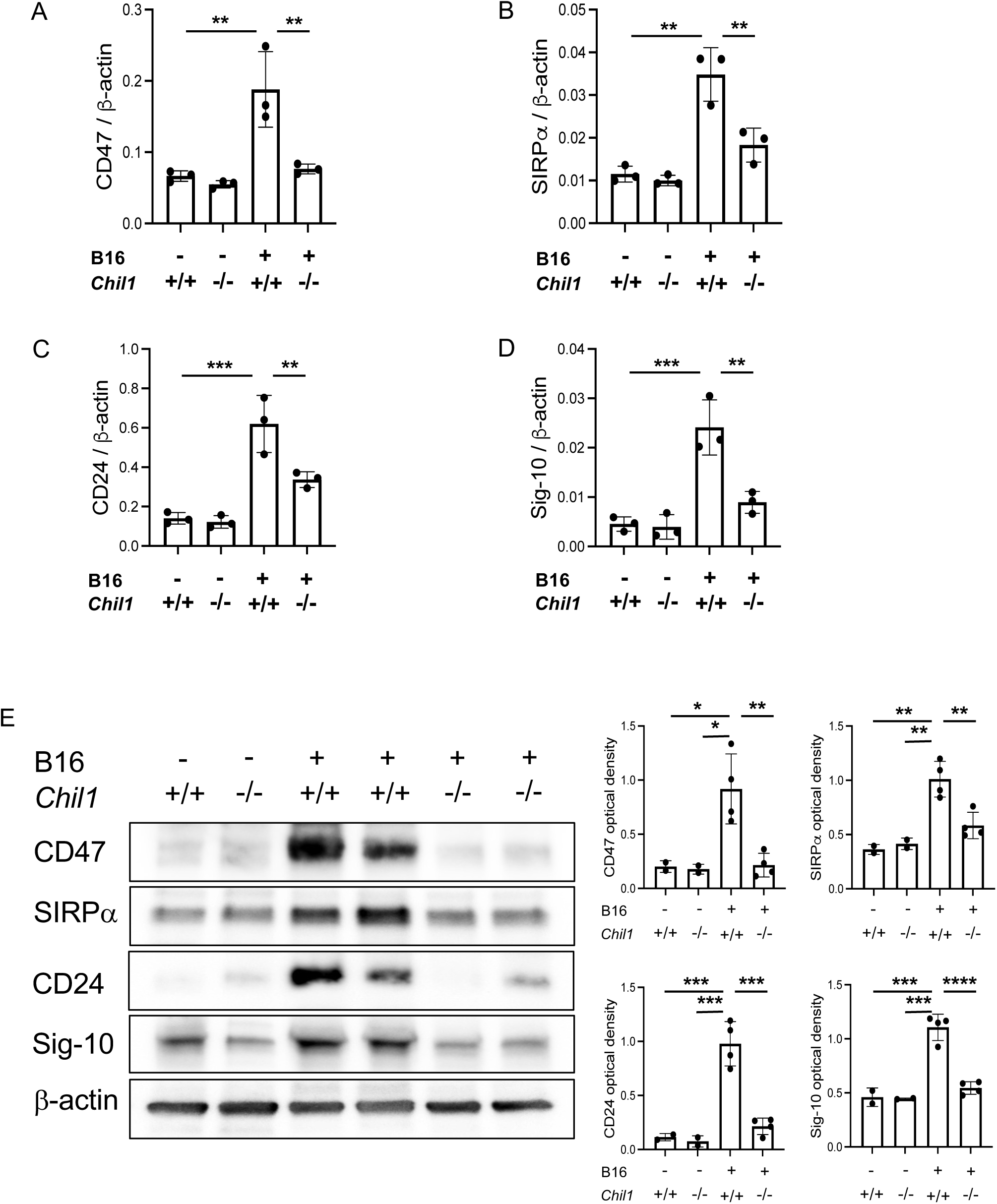
Pulmonary melanoma metastasis stimulates “don’t eat me” signals via a CHI3L1-dependent mechanism. (A-D) B16-F10 (B16) melanocytes were injected by tail vein into wild type mice (*Chil1*^+/+^) and Chi3l1 null (*Chil1*^-/-^) mice and the levels of whole lung mRNA encoding CD47 and its ligand SIRPα and CD24 and its ligand Siglec10 were quantitated 2 weeks later by qRT-PCR. (E) Lung lysates were also prepared, and Western blotting was used to quantitate the levels of CD47, SIRPα, CD24 and Siglec10 (Sig-10) proteins (left panel). The bar graphs in panel E represent the individual band intensity relative to β-actin control quantitated by densitometric analysis. Each dot in panels A-D represents an evaluation in a single mouse and the values in these panels represent the mean±SEM of these evaluations. Each dot in the densitometric evaluations represents an individual Western evaluation. \**P*<0.05, \*\**P*<0.01, \*\*\**P*<0.001, \*\*\*\**P*<0.0001.

### Anti-CHI3L1 abrogates melanoma stimulation of the CD47-SIRPα and CD24-Siglec10 “don’t eat me” phagocytic pathway(s)

To further define the role(s) of CHI3L1 in the stimulation of the CD47-SIRPα and CD24-Siglec10 pathways, we compared the expression of the moieties in wild type (WT) mice that had been randomized to receive B16 melanoma cells or controls and anti-CHI3L1 (FRG +) or its isotype control (FRG-) antibodies. As can be seen in Figure 2 (A-D), the stimulation of CD47, SIRPα, CD24 and Siglec10 by pulmonary metastasis was abrogated in mice that were treated with FRG. FRG had similar inhibitory effects on the levels of accumulation of CD47, SIRPα, CD24 and Siglec10 protein when assessed by Western blot evaluations (Figure 2E). Similar effects were seen with immunohistochemistry (IHC) (Supplemental Figure S1). When viewed in combination, these studies further demonstrate that anti-CHI3L1 is a powerful regulator of the induction of “don’t eat me” pathways during pulmonary metastasis.

**Figure 2.**
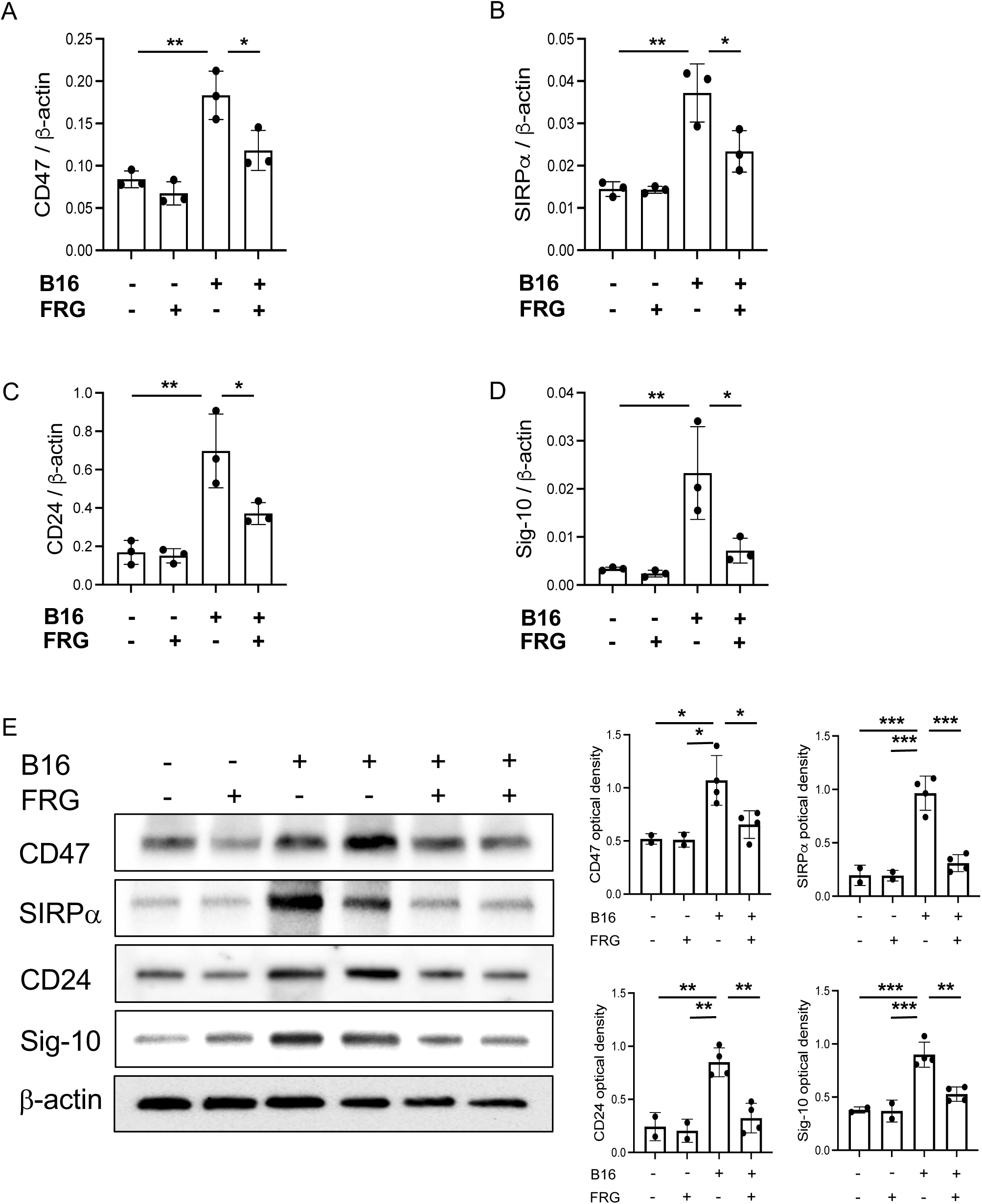
Anti-CHI3L1 ameliorates melanoma stimulation of “don’t eat me” signals. B16-F10 (B16) melanocytes were injected by tail vein into wild type mice and the mice were randomized to receive anti-CHI3L1 (FRG+) or its isotype control (IgG2b). (A-D) The levels of lung mRNA encoding CD47 and its ligand SIRP*α* and CD24 and its ligand Siglec10 were quantitated 2 weeks later. (E) Lung lysates were subjected to Western blotting to quantitate the levels of CD47, SIRPα, CD24 and Siglec10 (Sig-10) proteins. The bar graphs in panel E represent the individual band intensity relative to β-actin control quantitated by densitometric analysis. Each dot in panels A-D represents an evaluation in a single mouse and the values in these panels represent the mean±SEM of these evaluations. Each dot in the densitometric evaluations represents an individual Western evaluation. \**P*<0.05, \*\**P*<0.01, \*\*\**P*<0.001.

### Transgenic CHI3L1 stimulates the CD47-SIRPα and CD24-Siglec10 pathways in the absence of tumor cells

The studies noted above demonstrate that melanoma lung metastasis stimulates “don’t eat me” phagocytic pathways and that CHI3L1 plays an essential role in these regulatory events. Because a null mutation of CHI3L1 or treatment with a neutralizing anti-CHI3L1 antibody reduced metastatic spread (3, 4), they do not determine if the stimulatory effects are due to direct effects of CHI3L1 or the decreased metastatic spread of melanoma. To test this notion, we determined if CHI3L1 stimulates these phagocytosis inhibiting moieties without melanoma metastasis. In these experiments, the expression and accumulation of CD47 and its ligand SIRPα, and CD24 and its ligand Siglec10 were evaluated in lungs from WT mice and CHI3L1 overexpressing transgenic mice (*CHI3L1* Tg). These studies demonstrated that CHI3L1 itself is a potent stimulator of these moieties in lungs from *CHI3L1* Tg mice when compared to WT controls (Supplemental Figure S2). Thus, these studies demonstrate that the stimulation of CD47 and CD24 and their ligands in melanoma lung metastasis is mediated, at least in part, by tumor-independent effects of CHI3L1.

### CHI3L1 stimulates cell “don’t eat me” signals in vitro

CD47 is widely displayed on many, if not all cell types and SIRPα is characteristically found on myeloid cells (48). CD24 is a GPI anchored glycoprotein that is widely expressed on many cells in the immune system, frequently overexpressed in cancers and interacts with Siglec10 on macrophages (49). To further understand the regulation of “don’t eat me signals”, we characterized the effects of rCHI3L1 on B16 melanocytes and RAW macrophages. These studies demonstrated that CHI3L1 is a potent stimulator of CD47, CD24 and Siglec10 but not SIRPα mRNA and protein in B16 cells (Figure 3, A and B). They also demonstrated that CHI3L1 also stimulates CD47, SIRPα, CD24 and Siglec10 mRNA and protein in RAW macrophages (Figure 3, C and D).

**Figure 3.**
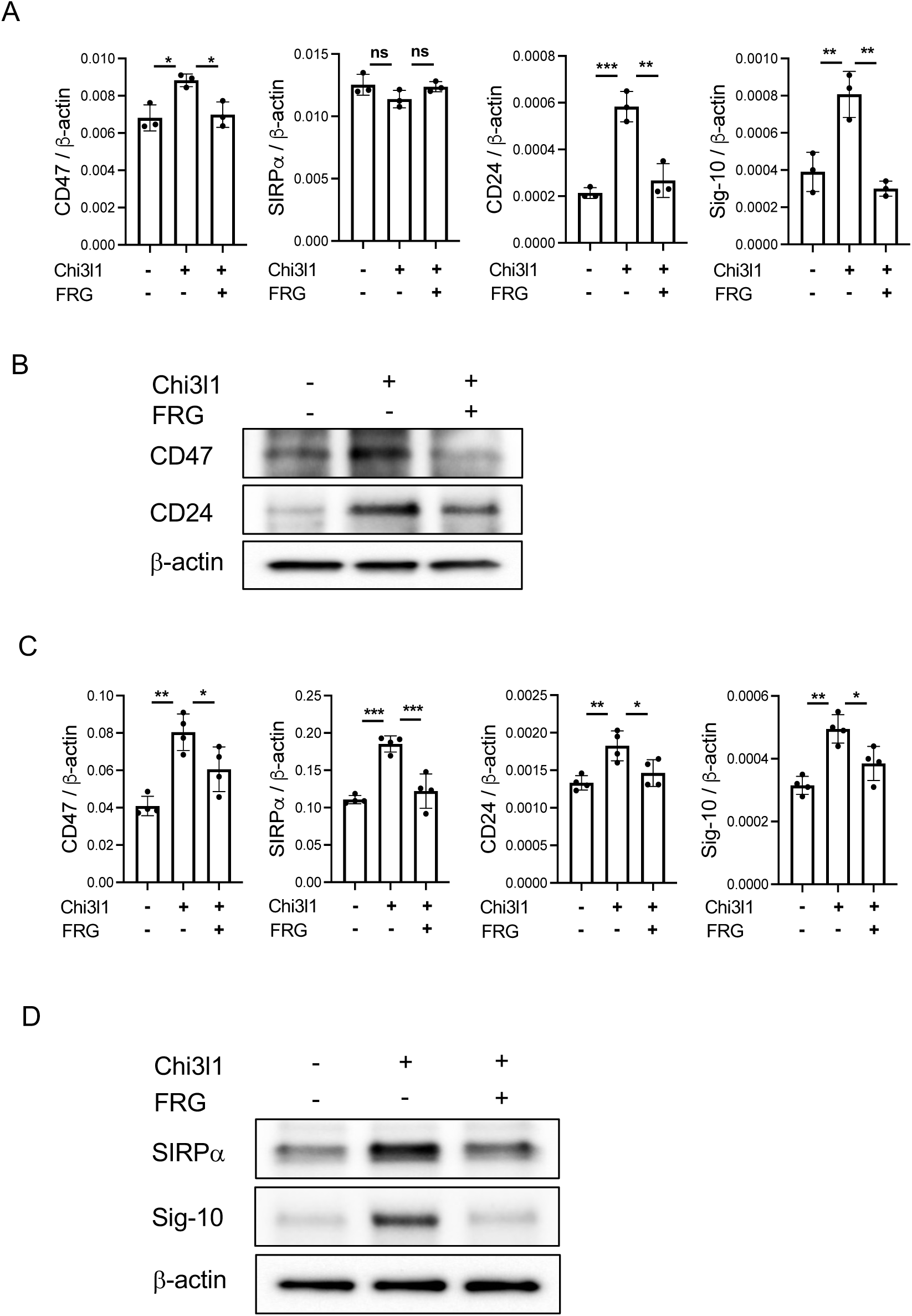
CHI3L1 regulation of melanocyte and RAW macrophage “don’t eat me” signals. (A and B) B16-F10 melanoma cells were incubated with rCHI3L1 alone or together with anti-CHI3L1 FRG antibody and the levels of CD47, SIRPα, CD24 and Siglec10 mRNA (A) and protein (B) were evaluated. (C and D) RAW cells were incubated with rCHI3L1 alone or together with anti-CHI3L1 FRG antibody and the levels of CD47, SIRPα, CD24 and Siglec10 mRNA (C) and protein (D) were evaluated. The bar graphs in panels B and D represent the individual band intensity relative to β-actin control quantitated by densitometric analysis. In panels A and C, each dot represents the evaluation in an individual animal and the values in these panels represent the mean±SEM of these evaluations. Panels B and D are representative of a minimum of 3 similar evaluations. \**P*<0.05; **P<0.01; \*\*\**P*<0.001, ns=not significant.

### Pulmonary melanoma metastasis inhibits “eat me” phagocytic pathways via a CHI3L1-dependent mechanism

Studies were next undertaken to determine if major “eat me” phagocytic pathways were regulated by pulmonary metastasis. In these experiments wild type C57BL/6 mice were challenged with freshly prepared B16 melanoma cells (B16+) or vehicle control (B16-) and the levels of mRNA encoding calreticulin and HMGB1 were assessed 2 weeks later. These studies demonstrated that B16 melanoma metastasis decreased the levels of expression of these moieties (Figure 4, A and B). Comparable alterations in calreticulin and HMGB1 protein accumulation were seen in Western evaluations (Figure 4C). These inhibitory effects were mediated, at least in part, by CHI3L1 because CHI3L1 inhibition of these “eat me” signals was significantly restored in *Chil1* null (*Chil1*^-/-^) mice (Figure 4, A-C). Similarly, treatment with FRG (FRG+) diminished the inhibition of “eat me” signal mRNA and protein when compared to isotype controls (FRG-) (Figure 4, D-F). When viewed in combination with the studies noted above, these studies demonstrate that melanoma lung metastasis stimulates the expression of “don’t eat me” and diminishes the expression and accumulation of “eat me” phagocytic signals via CHI3L1-dependent mechanisms.

**Figure 4.**
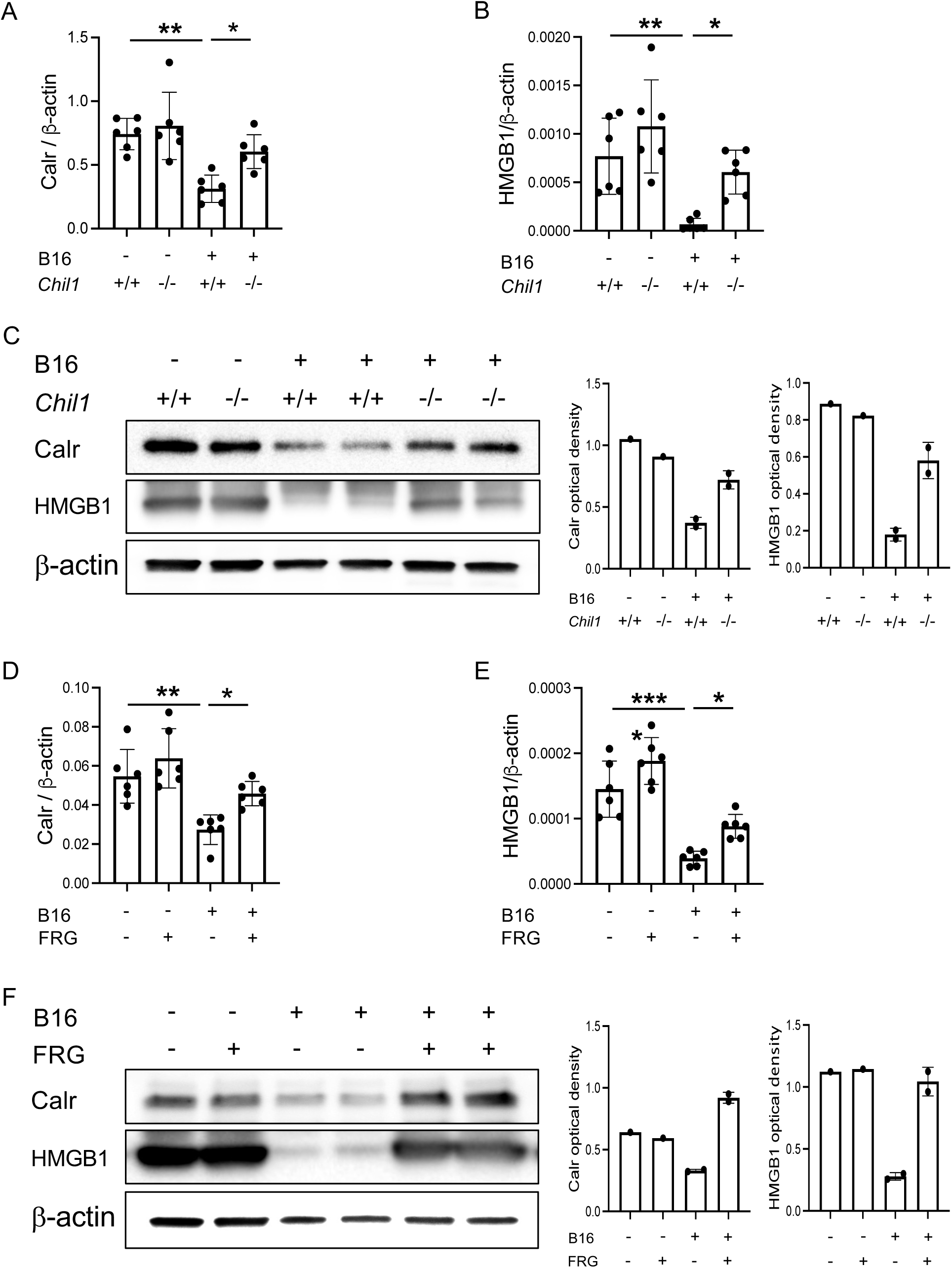
Metastasis and anti-CHI3L1 regulation of “eat me” signals. B16-F10 (B16) melanocytes were injected via tail vein into wild type mice (*Chil1*^+/+^) and Chi3l1 null (*Chil1*^-/-^) mice. (A-C) The levels of lung calreticulin (Calr) and HMGB1 mRNA (A and B) and protein (C) were quantitated 2 weeks later. (D-F) B16-F10 melanocytes were also injected through tail vein into wild type mice that were randomized to receive FRG (FRG+) or its isotype control antibody (FRG -). The levels of lung *HMGB1* or *Calr* mRNA (D and E) and protein expression (F) were assessed 2 weeks later. In panels A, B, D, and E, each dot represents the evaluation in an individual animal and the values in these panels represent the mean±SEM of these evaluations. The densitometric bar graphs in panels C and F represent the individual band intensity of the noted experiment relative to the β-actin control quantitated by densitometric analysis. \**P*<0.05; \*\**P*<0.01; \*\*\**P*<0.001, \*\*\*\**P*<0.0001.

### Recombinant CHI3L1 inhibits murine macrophage phagocytosis

To further understand the effects of CHI3L1 on macrophage phagocytosis we incubated murine macrophages with rCHI3L1 or vehicle control and evaluated their ability to ingest zymosan particles using fluorescence microscopy. As can be seen in figure 5A, RAW macrophages incubated with vehicle control readily ingested the green zymosan fluorescent bioparticles. In contrast, incubation with rCHI3L1 markedly inhibited these phagocytic responses (Figure 5A). These inhibitory alterations were direct effects of rCHI3L1 because the ability of these cells to ingest zymosan was restored when the cells were treated with FRG (compared to its Isotype control) or the small molecule CHI3L1 inhibitor kasugamycin (Figure 5A). To be sure that these findings reflected true phagocytosis, cells were also evaluated using pHrodo Red-633 which emits a yellow signal when ingested into a low pH environment such as a phagocytic compartment. As noted with zymosan, these experiments demonstrated that CHI3L1 inhibited RAW cell ingestion of pHrodo Red when compared to cells incubated in vehicle control and this inhibition was abrogated by treatment with FRG or kasugamycin (Figure 5B). The magnitude of these effects of CHI3L1 can be seen with FACS evaluations which demonstrated that rCHI3L1 was a powerful inhibitor of macrophage phagocytosis and that these effects were abrogated by treatment with FRG or kasugamycin (Figure 5C).

**Figure 5.**
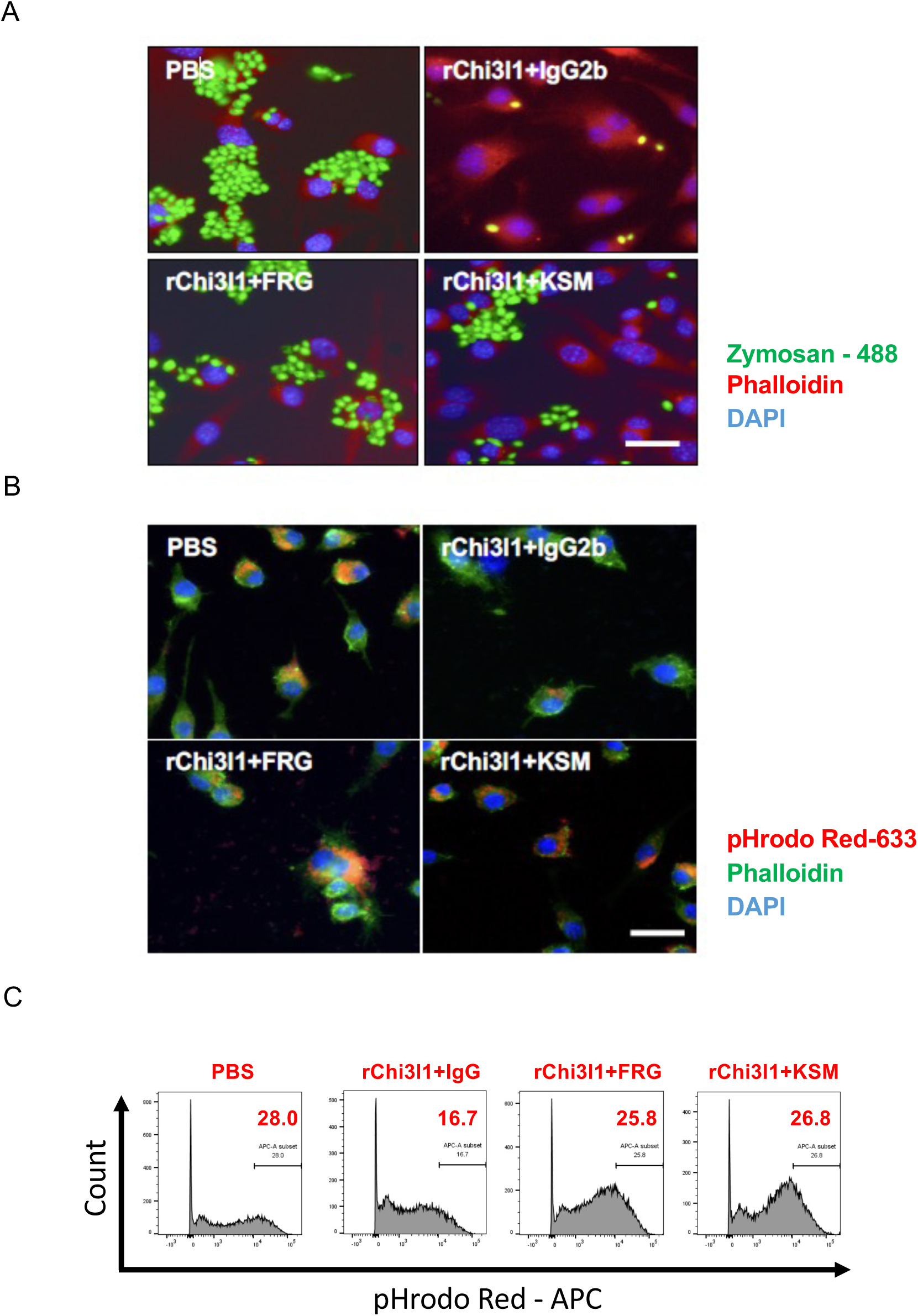
CHI3L1 regulation of murine macrophage phagocytosis. (A) RAW cells were incubated with zymosan-488 in the presence of anti-CHI3L1 (FRG), its isotype control (IgG2b) or Kasugamycin (KSM). Nuclei are highlighted with blue (DAPI) staining and cellular structural motifs are stained red with phalloidin. (B) Cells were also evaluated using pHrodo Red-633 which emits a yellow signal when ingested into a low pH environment such as a phagocytic compartment. (C) FACS was also used to evaluate the levels of RAW cell phagocytosis of zymosan-488. These experiments compared zymosan ingestion in the presence of rCHI3L1 or its vehicle control and in the presence of FRG, kasugamycin or their controls. A-C are representative of a minimum of 3 similar evaluations. In panel A and B, scale bars=25μm. The scale bar in lower right panel applies to all the other panels.

### Recombinant CHI3L1 inhibits human macrophage phagocytosis

To be sure that the effects of rCHI3L1 noted above were not murine-specific, similar evaluations were undertaken with human THP-1 cells. These cells were initially incubated with PMA and then rCHI3L1 to induce M0 and then M2 differentiation as described by our laboratory and others (33, 50). This was followed by exposure to zymosan and immunofluorescent evaluation of particle ingestion. In accord with the findings with RAW cells, the levels of mRNA encoding the “don’t eat me” signals CD47, SIRPα, CD24 and Siglec10 were increased by rCHI3L1 in PMA pretreated THP-1 cells (Figure 6, A-D) Similarly, the PMA pretreated THP-1 cells manifest impressive levels of zymosan phagocytosis that was markedly decreased after incubation with rCHI3L1 (Figure 6E). Importantly, rCHI3L1 inhibition of phagocytosis was restored by treatment with FRG or kasugamycin (Figure 6, A-E). When viewed in combination with the findings noted above, these studies demonstrate that CHI3L1 directly inhibits phagocytosis while inducing “don’t eat me” signals in human as well as murine macrophages.

**Figure 6.**
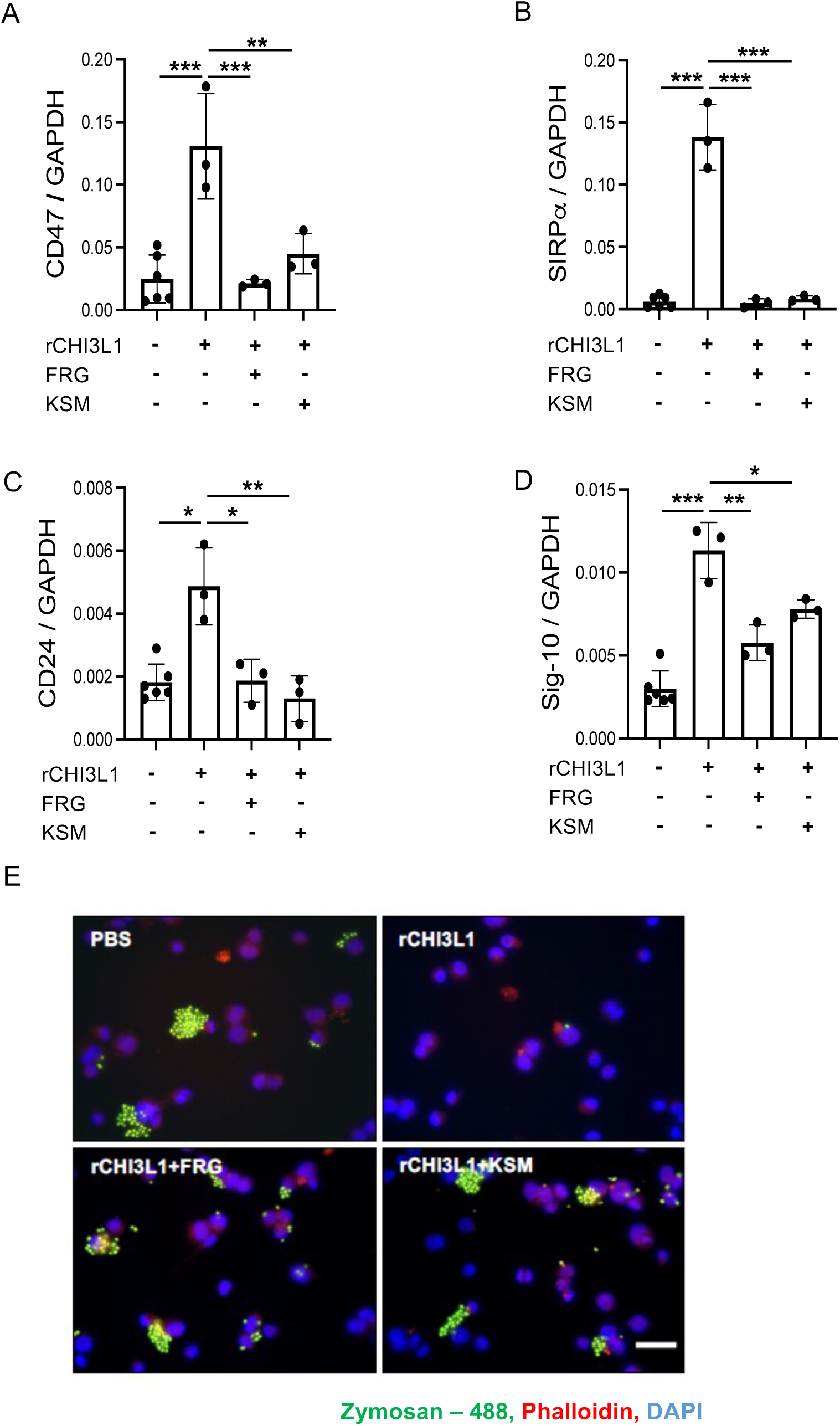
CHI3L1 regulation of PMA-treated human THP-1 cell expression of “don’t eat me” signals and zymosan bioparticle phagocytosis. Human THP-1 cells were differentiated into M0 cells by treatment with PMA. They were then incubated with rCHI3L1 or vehicle control in the presence (+) or absence (−) of FRG or kasugamycin. (A-D) The levels of mRNA encoding CD47, SIRPα, CD24 and Siglec10 were evaluated by qRT-PCR. (E) Zymosan phagocytosis analysis. Each dot in panels A-D represents the evaluation in an individual *in vitro* experiment. Panel E is representative of 3 similar evaluations \**P*<0.05; \*\**P*<0.01, \*\*\**P*<0.001. In panel E, scale bar=25μm. The scale bar in lower right panel applies to all the other panels.

### Pulmonary melanoma metastasis stimulates SHP-1 and SHP-2 via a CHI3L1-dependent mechanism

The binding of CD47 and SIRPα induces the protein tyrosine phosphatases SHP-1 and SHP-2 (14). Thus, studies were undertaken to determine if these inductive events were altered during pulmonary melanoma metastasis and if CHI3L1 plays an important role in these regulatory events. In these experiments, C57BL/6 mice were challenged with freshly prepared B16-F10 melanoma cells or vehicle control and the levels of lung mRNA encoding SHP-1 and SHP-2 were assessed 2 weeks later. These studies demonstrated that B16 melanoma metastasis stimulated the levels of expression of both of these phosphatase moieties (Figure 7, A and B). Comparable alterations in SHP-1 and SHP-2 protein accumulation were seen in Western evaluations (Figure 7C). These inhibitory effects were mediated, at least in part, by CHI3L1 because the induction of SHP1 and SHP-2 were markedly decreased in *Chil1* null (*Chil1*^-/-^) mice and wild type mice treated with FRG versus an isotype control (Figure. 7, A-F). These studies demonstrate that melanoma lung metastasis, while stimulating the CD47-SIRPα and CD24-Siglec10 phagocytosis inhibiting pathways, also induce SHP-1 and SHP-2 via a CHI3L1-dependent mechanism(s).

**Figure 7.**
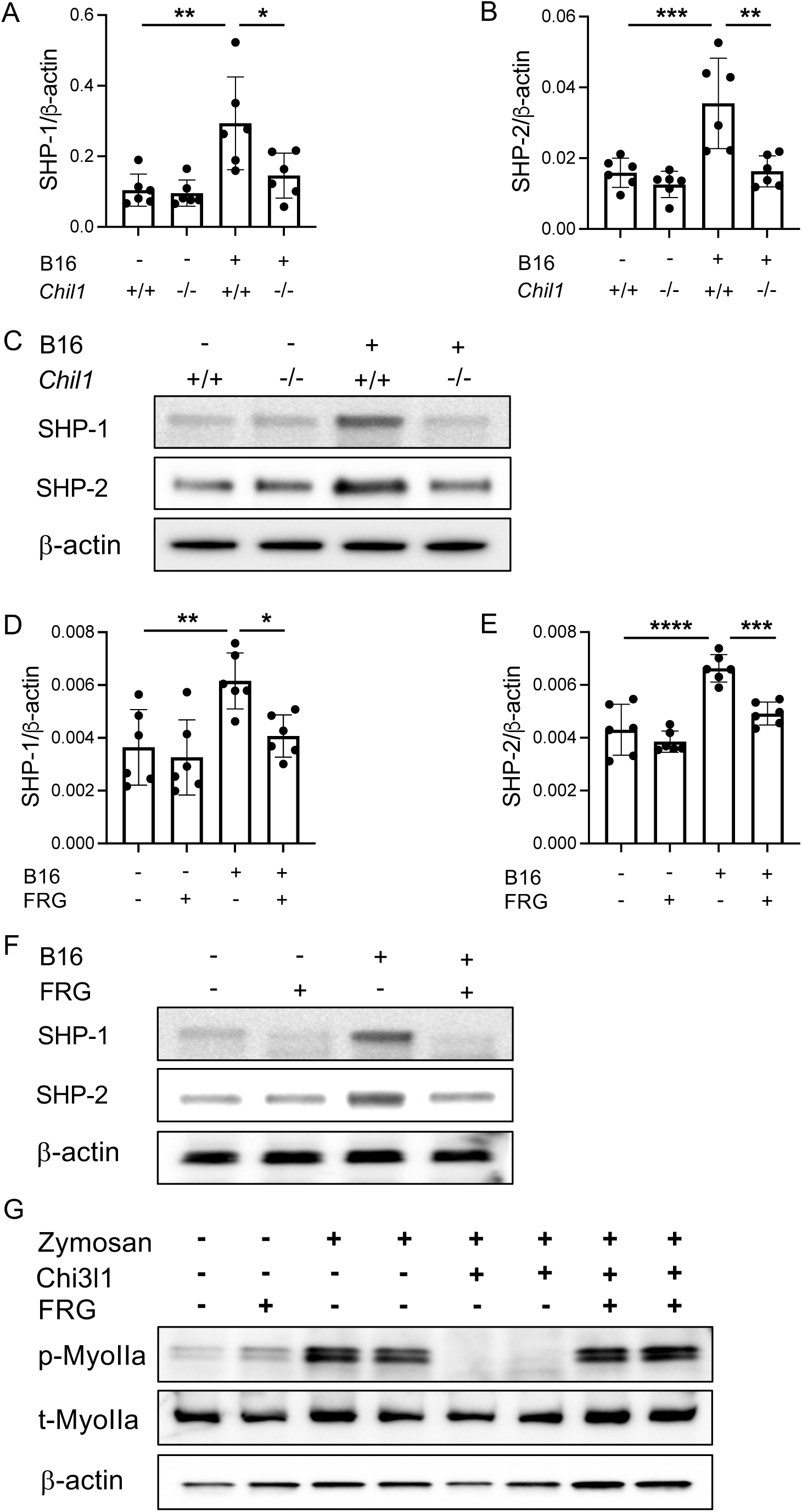
CHI3L1 regulation of SHP-1 and SHP-2 mRNA and protein *in vivo* and RAW cell Myosin IIa *in vitro*. (A and B) B16-F10 melanocytes were injected by tail vein into wild type mice (*Chil1^+/+^*) and Chi3l1 null (*Chil1^-/-^)* mice and the levels of whole lung mRNA encoding SHP-1 and SHP-2 were quantitated 2 weeks later. (C) Lung lysates were also prepared and Western blotting was used to quantitate the levels of SHP-1 and SHP-2 proteins. (D-F) Regulation of SHP-1 and SHP-2 in mice randomized to receive FRG or its isotype control. (G) RAW cells were also incubated with zymosan, recombinant rCHI3L1 and FRG or appropriate controls and the levels of total (t) or phosphorylated (p) non-muscle Myosin IIa (NMIIa) were assessed using Western blots evaluations. The densitometric bar graphs in panels C, F, and G represent the individual band intensity of the noted experiment relative to the β-actin control quantitated by densitometric analysis. Each dot in panels A, B, D and E represents an evaluation in a single mouse. Panels C, F and G are representative of at least 3 similar evaluations. \**P*<0.05; \*\**P*<0.01; \*\*\**P*<0.001, \*\*\*\**P*<0.0001.

### CHI3L1 inhibits the expression and the phosphorylation of non-muscle Myosin IIa

To gain insight into the mechanism(s) that CHI3L1 uses to regulate phagocytosis, we evaluated the effects of CHI3L1 on macrophage non-muscle myosin IIa (NMIIa). At baseline, total (t) NMIIa and, to a lesser extent, phosphorylated (p)-NMIIa were appreciated at modest levels in RAW cells (Figure 7G). Zymosan phagocytosis caused a marked increase in the levels of p-Myosin protein accumulation and a modest increase in total-Myosin IIa (Figure 7G). Importantly, treatment with rCHI3L1 caused a modest decrease in t-NMIIa and a marked decrease in p-NMIIa (Figure 7G). This effect was ameliorated, t-NMIIa was modestly increased and p-NMIIa accumulation was markedly augmented when the cells were treated with FRG compared to isotype controls (Figure 7G). These studies demonstrate that CHI3L1 inhibition of phagocytosis is associated with a modest decrease in NMIIa accumulation and a major decrease in NMIIa phosphorylation.

## Discussion

Cancer immunotherapy began in 1893 with Coley’s toxin (14). This was followed, in the 1950s, by the immune surveillance hypothesis that proposed that the immune system could recognize and reject cancer cells as foreign and attack and kill them (51, 52). These seminal observations have opened the door to an impressive body of research highlighting the importance of anti-cancer immune responses. The majority of these studies have focused on adaptive immune responses and T cells, particularly CD8+ cells. The success of this focus is readily apparent with interventions that target ICPs such as PD-1, PD-L1 and CTLA4 manifesting impressive therapeutic efficacy in a variety of tumors (53). Surprisingly, although innate immune responses play critical roles in the regulation of cancer initiation and progression, and are a prerequisite for adaptive anti-tumor immune responses, the importance of innate immune events such as phagocytosis and the mononuclear cells that mediate these responses have not been appropriately evaluated. Previous studies from our laboratory demonstrated that CHI3L1 is induced during neoplasia and highlighted its impressive ability to inhibit adaptive immune responses via its ability to stimulate ICPs pathways and inhibit T cell costimulation (4, 54). However, the mechanisms that mediate these responses have not been adequately defined. In particular, although phagocytosis plays a critical role in T cell antigen priming and the generation of adaptive immunity, the importance of CHI3L1 as a regulator of innate immunity and phagocytosis have not been adequately defined.

To further understand the roles of CHI3L1 and innate immunity in anti-tumor responses we characterized the effects of CHI3L1 on mononuclear cell phagocytosis and the pathways that regulate these responses. These studies highlight the ability of pulmonary metastasis to stimulate “don’t eat me” phagocytic pathways including the CD47-SIRPα pathway and the CD24-Siglec10 pathway while inhibiting “eat me” signals from moieties such as calreticulin and HMGB1. They also demonstrate that these events are stimulated by CHI3L1, highlight the ability of CHI3L1 to directly inhibit phagocytosis and define the role(s) of Myosin IIa in mediating these regulatory events. Lastly, these findings have important therapeutic implications. By demonstrating that interventions that block CHI3L1 inhibit innate antitumor responses they define a mechanism that CHI3L1 uses in its role as a master regulator of ICPs and adaptive antitumor responses. They also reinforce the import of anti-CHI3L1-based cancer therapies as interventions that simultaneously regulate adaptive and innate anti-tumor responses.

Cancer cells try to evade phagocytic clearance, the expression of neoantigens with MHC class I on the cell surface and the development of adaptive immune responses via the augmented expression of antiphagocytic ligands (14, 15). One of the earliest phagocytic checkpoints to be recognized was CD47 which is a transmembrane protein that is broadly expressed on normal cells confirming their recognition as “self” and inhibiting their phagocytic ingestion. It is also induced in and expressed on tumor cells and was shown to bind to its ligand SIRPα on macrophages (14, 55). In contrast to CD47, SIRPα expression is limited to myeloid cells and neutrophils (14, 56). This binding triggers the SIRPα immunoreceptor tyrosine based inhibitory motifs (ITIMs) which activate the SHP-1 and SHP-2 phosphatases which dephosphorylate myosin IIa and inhibit phagocytosis (14, 57). Our studies add to our existing understanding of the CD47-SIRPα axis by demonstrating that melanoma metastasis stimulates CD47 and SIRPα on epithelial cells, macrophages and tumor cells. They also demonstrate that these events are mediated by CHI3L1 which also stimulates SHP-1 and SHP-2 and disrupts and dephosphorylates myosin IIa. In keeping with the “don’t eat me “signal that CD47-SIRPα binding conveys, our studies also demonstrate that CHI3L1 directly inhibits murine and human macrophage phagocytosis. Because macrophages play a critical role in antigen processing priming of T cells, one can see how the induction of these “don’t eat me” signals inhibit phagocytosis and diminishes adaptive immune responses against tumor neoantigens. In addition, because the expression of CD47 on tumors correlates inversely with patient prognosis (14, 58–61), one can see how interventions that inhibit the interaction of CD47 and SIRPα such as treatment with anti-CHI3L1, have impressive potential as cancer therapeutics.

CD24 is a heavily glycosylated GPI-anchored surface protein that interacts with Siglec10 on immune cells in order to dampen a wide range of damaging inflammatory responses (49). Unlike CD47 which is widely expressed on normal cells, CD24 is also expressed on tumor cells where it is thought of as a tumor-specific marker (49, 62, 63). Siglec10 is stimulated by the inhibitory cytokines IL-10 and TGF-β1 and the Th2 cytokine IL-4. In all cases, CD24-Siglec10 binding elicits an inhibitory signaling cascade mediated by SHP-1 and SHP-2 which blocks innate immune responses and phagocytosis (49). Our studies add to our existing knowledge of the regulation of “don’t eat me” signals by demonstrating that CHI3L1 is a critical stimulator of CD24 and Siglec10 and a potent stimulator of SHP-1 and SHP-2 phosphatases (49). It is also important to point out that, in most cases, CD24 blockade by itself potentiates phagocytosis and that dual CD24 and CD47 blockade augment phagocytosis at least additively (49). Collectively, our studies support the concept that CD24 is a novel innate immune checkpoint that plays a critical role in antitumor immunity and that CD24 inhibitors have impressive therapeutic potential. Our studies also add to this vision by demonstrating that CHI3L1 simultaneously augments the CD24-Siglec10 and the CD47-SIRPα phagocytosis inhibiting pathways. This provides additional support for the potential importance of anti-CHI3L1-based interventions that block both of these pathways in the treatment of cancer and other disorders.

Macrophages are immune cells with impressive phenotypic heterogeneity that are indispensable in a variety of biologic responses including organ development, tissue turnover, inflammation, tumorigenesis and metabolism (64). Studies of this heterogeneity have revealed complex mechanisms of macrophage activation with M0 cells manifesting the ability to differentiate into opposing M1 and M2 polarization states (65). They also demonstrated that macrophages play important roles in tumorigenesis, highlighting the tumor-stimulating effects of M2 or M2-like TAMs (tumor associated macrophages) and high antigen presenting capacity and augmented antitumor effects of M1 TAM (64). Previous studies from our laboratory and others demonstrated that CHI3L1 is a powerful stimulator of M2 macrophage differentiation (23). In accord with these findings, the present studies demonstrate that PMA stimulated human THP-1 cells, which have a M0 phenotype, take on a M2 phenotype when incubated with CHI3L1. They also add to our understanding of this response by demonstrating that CHI3L1 is a potent inhibitor of phagocytosis in murine and human cell systems. This is in keeping with the known ability of the TH2 cytokine IL-4 to stimulate the phagocytosis inhibitor Siglec10 (49). When viewed in combination, these studies demonstrate that M2 macrophages have a diminished phagocytic capacity and a diminished ability to ligate the innate and adaptive immune responses when compared to M0 and M1 macrophages (14, 64).

Cell death and cellular turnover are fundamental to healthy living (1). Roughly 200 billion cells are turned over in the human body daily via apoptosis and the coordinated daily removal of these apoptotic cells, termed efferocytosis (1). This apoptotic process and the removal of apoptotic cells is critical for routine tissue homeostasis, embryonic development, wound healing, and inflammation resolution (66, 67). Failed or defective clearance of apoptotic cells can be causative in numerous pathologies including atherosclerosis, aging-associated inflammation, cancer and inflammation (68). Dysfunctional efferocytosis is also seen in COPD and pulmonary fibrosis (7, 69, 70). COPD is characterized by the accumulation of uncleared apoptotic cells that may undergo secondary necrosis. Deficiencies in efferocytosis in COPD can persist for years and cause pulmonary function failure and tissue destruction (71). Idiopathic Pulmonary Fibrosis (IPF) is characterized by a sustained epithelial insult(s) with dysregulated fibroproliferative repair (72). It is also characterized by the accumulation of apoptotic cells and a reduced rate of efferocytosis by alveolar macrophages which leads to fibrosis (69). It has also been reported in an animal model of pulmonary fibrosis that efferocytosis attenuates lung injury by inducing the production of hepatocyte growth factor (HGF) and driving fibroproliferative repair (73). Interestingly, the levels of CHI3L1 are increased in COPD and IPF. One can see how the ability of CHI3L1 to stimulate “don’t eat me” and inhibit “eat me” signals can inhibit phagocytosis and cause dysfunctional efferocytosis and its inflammatory and fibrotic tissue consequences. One can also see how this suppressed phagocytic response can lead to the defective adaptive immune responses that have been described in COPD (47) and other disorders (7) and how anti-CHI3L1 based interventions can be therapeutically effective in these diseases.

A fundamental challenge in cancer therapy and biology relates to the appreciation that successful interventions are often transient and are thwarted by the development of therapeutic resistance. To address this issue, therapeutic regimens are often designed to simultaneously focus on multiple oncogenic pathways based on the assumption that while resistance to one pathway may develop, resistance to multiple pathways is less likely to occur. The present studies demonstrate that CHI3L1 inhibits innate immune responses via inhibiting phagocytosis. Prior studies from our laboratory and others have also demonstrated that CHI3L1 is a master regulator of ICPI and T cell co-stimulation, and an important regulator of B-raf, p53, PTEN, and other tumor suppressors (3, 4, 74). Taken together, these studies demonstrate that CHI3L1-based interventions are particularly intriguing as cancer therapeutics because they simultaneously control multiple oncogenic pathways.

Phagocytic cells ingest cells or particles via a coordinated process of adhesion, pseudopod extension and internalization and phagosome closure (57). It is initiated by the activation of cytoskeletal assembly by non-muscle myosin, myosin IIa (NMIIa), which accumulate and act as contractile motors during particle/cell internalization (57). A key component of this process is how the macrophage differentiates foreign cells and or particles from “self”. As noted above, the ubiquitous expression of CD47 suggests that it is an important phagocytosis inhibiting “marker of self” (57). The present studies add to our understanding of the phagocytic process by demonstrating that NMIIa is the likely contractile motor driving cell/particle engulfment. Importantly, they also demonstrate that CHI3L1 is a powerful inhibitor of NMIIa accumulation and phosphorylation and, as a consequence, cell/particle engulfment and overall phagocytosis. These findings are in accord with prior observations that demonstrated that NMIIa is a direct target of SHP-1 and that CD47 selectively downregulates cytoskeletal contributions to phagocytic engulfment (57).

In conclusion, these studies demonstrate that CHI3L1 inhibits macrophage phagocytosis by stimulating the CD47-SIRPα and CD24-Siglec10 phagocytosis checkpoint pathways while inhibiting macrophage “eat me” signals including those provided by calreticulin and HMGB1. This inhibition of innate immunity provides a mechanistic explanation for the ability of CHI3L1 to stimulate ICPI and inhibit adaptive immune responses in cancer and diseases like COPD. The link between the ability of CHI3L1 to inhibit innate and adaptive immune responses provides additional insight into the many oncogenic, inflammatory and remodeling responses mediated by CHI3L1 and provides rationale for the use of CHI3L1-based interventions in the treatment of cancer and other disorders. Additional investigations of the biology and therapeutic implications of CHI3L1 regulation of innate immunity and other pathways are warranted.

