## Supplemental data and figures for "Chitinase 3-like-1 (CHI3L1) Inhibits Innate Anti-Tumor and Tissue Remodeling Immune Responses by Regulating CD47-SIRPα and CD24-Siglec10-Mediated Phagocytosis"

#### Supplementary Table

##### Table S1 Sequences of RT-PCR primers used in this study

| Genes | Sequences (5' to 3') |
| --- | --- |
| CD47-S | TGGTGGGAAACTACACTTGCG |
| CD47-AS | CGTGCGGTTTTTCAGCTCTAT |
| SIRPa-S | CCACGGGGAAGGAACTGAAG |
| SIRPa-AS | ACGTATTCTCCTGCGAAACTGTA |
| CD24-S | GTTGCACCGTTTCCCGGTAA |
| CD24-AS | CCCCTCTGGTGGTAGCGTTA |
| Sigl-G-S | GTCCCAGACTTGCATGAGAATC |
| Sigl-G-AS | GACCCAGCTCAGTGTAGCA |
| Calr-S | GCAGACCCTGCCATCTATTTC |
| Calr-AS | TCGGACTTATGTTTGGATTCGAC |
| HMGB1-S | GGCGAGCATCCTGGCTTATC |
| HMGB1-AS | GGCTGCTTGTCATCTGCTG |
| SHP1-S | GGACTTCTATGACCTGTACGGA |
| SHP1-AS | GCTGCGTGTAATACTCGACCA |
| SHP2-S | AGAGGGAAGAGCAAATGTGTCA |
| SHP2-AS | CTGTGTTTCCTTGTCCGACCT |

### Supplementary Figures

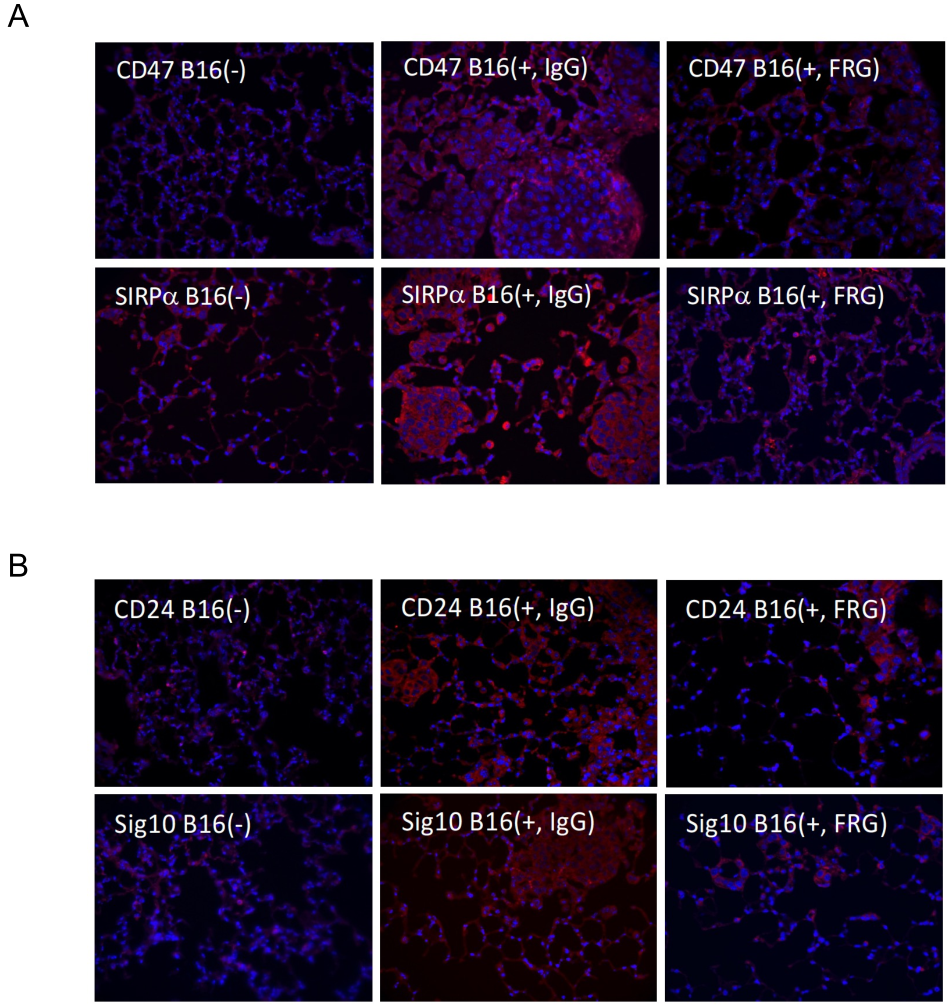


**Figure S1**. Expression of proteins associated with "don't eat me" phagocytic pathway in lungs with and without B16-F10 (B16) melanoma metastasis. B16-F10 melanocytes (B16(+)) or vehicle control (B16(-)) were injected by tail vein into wild type mice (*Chil1*+/+) and the mice were randomized to receive antiCHI3L1 (FRG+) or its isotype control (IgG2b). Sectioned lungs were stained with fluorescent labeled antibodies (red) against CD47, SIRPα, CD24, and siglec10 (Sig10). x40 of original magnification.

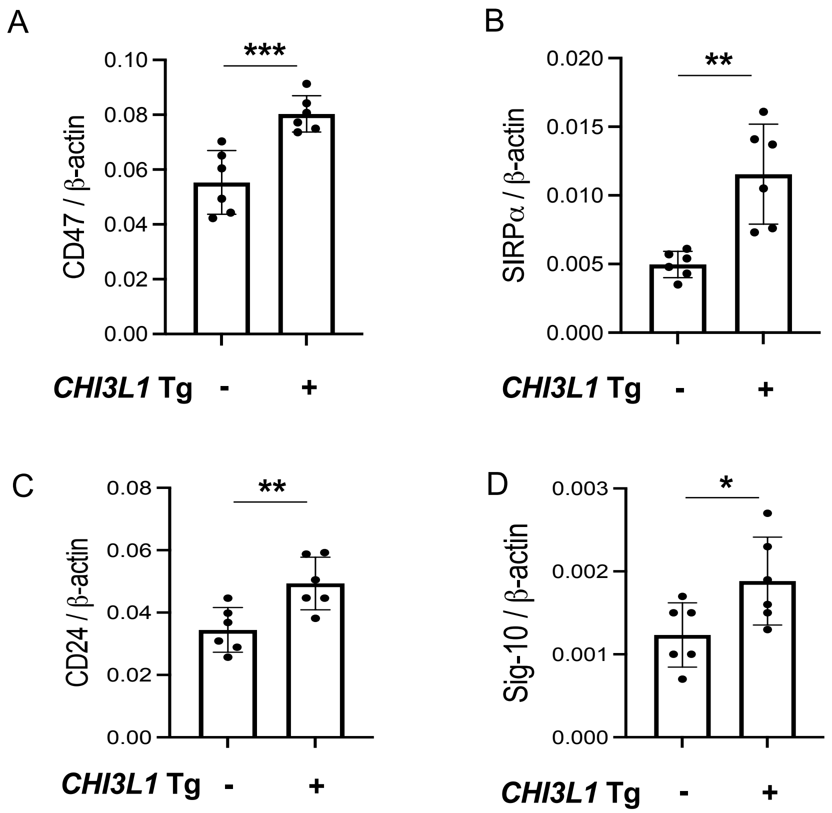


**Figure S2**. Expression of genes associated with “don’t eat me” phagocytic pathway in lungs from CHI3L1 transgenic (Tg+) mice and mtransgene negative (Tg-) controls. The levels of mRNA encoding CD47, SIRPα, CD24, and siglec10 (Sig-10) were detected by real-time RT-PCR (qRT-PCR). Each dot represents the evaluations in lungs from a single animal. *p<0.05, **P<0.01, ***P<0.001
